# A biochemical mechanism for time-encoding memory formation within individual synapses of Purkinje cells

**DOI:** 10.1101/596148

**Authors:** Ayush Mandwal, Javier G. Orlandi, Christoph Simon, Jörn Davidsen

## Abstract

Within the classical eye-blink conditioning, Purkinje cells within the cerebellum are known to suppress their tonic firing rates for a well defined time period in response to the conditional stimulus after training. The temporal profile of the drop in tonic firing rate, i.e., the onset and the duration, depend upon the time interval between the onsets of the training conditional and unconditional stimulus. Direct stimulation of parallel fibers and climbing fiber by electrodes was found to be sufficient to reproduce the same characteristic drop in the firing rate of the Purkinje cell. In addition, the specific metabotropic glutamate-based receptor type 7 (mGluR_7_), which resides on the Purkinje cell synapses, was found responsible for the initiation of the response, suggesting an intrinsic mechanism within the Purkinje cell for the temporal learning. In an attempt to look for a mechanism for time-encoding memory formation within individual Purkinje cells, we propose a biochemical mechanism based on recent experimental findings. The proposed model tries to answer key aspects of the “Coding problem” of Neuroscience by focussing on the Purkinje cell’s ability to encode time intervals through training. According to the proposed mechanism, the time memory is encoded within the dynamics of a set of proteins — mGluR_7_, G-protein, G-protein coupled Inward Rectifier Potassium ion channel, Protein Kinase A and Protein Phosphatase 1 — which self-organize themselves into a protein complex. The intrinsic dynamics of these protein complexes can differ and thus can encode different time durations. Based on their amount and their collective dynamics within individual synapses, the Purkinje cell is able to suppress its own tonic firing rate for a specific time interval. The time memory is encoded within the effective rate constants of the biochemical reactions and altering these rates constants means storing a different time memory. The proposed mechanism is verified by a simplified mathematical model and corresponding dynamical simulations of the involved biomolecules, yielding testable experimental predictions.

**Author summary:** Hebbian plasticity is a widely accepted form of learning that can encode memories in our brain. Spike-timing dependent plasticity resulting in Long-term Potentiation or Depression of synapses has become the default mechanistic explanation behind memory formation within a neuronal population. However, recent experiments of conditional eyeblink response in Purkinje cells have challenged this point of view by showing that these mechanisms alone cannot account for time memory formation in the Purkinje cell. To explain the underlying mechanism behind this novel synaptic plasticity, we introduce a biochemical mechanism based on protein interactions occurring within a single synapse. These protein interactions and the associated effective rate constants are sufficient to encode time delays by auto-induced inhibition on a single excitatory synapse, suggesting that synapses are capable of storing more information than previously thought.

## Introduction

How do we store memories in our brain? How do we retrieve and edit them when required? Recent experimental findings have shed some light onto these fundamental questions. Experiments have shown that memories are held within specific neuronal populations [1–3]. Such populations, referred as *memory engram cells* [4, 5] store memory either by forming or eliminating synapses [6, 7] or by altering synaptic strengths between neurons [8, 9] within the population. These forms of learning and memory formation fall under the widely accepted Hebbian learning paradigm [10]. However, the individual contribution of each synapse to the engrams, and how changes in synaptic strength affects memories, remain poorly understood. The problem of information encoding was raised by C.R. Gallistel [11] and termed as the “Coding Question”, one of the fundamental open questions in Neuroscience today. Recent experiments on Purkinje cells, one of the major neuronal populations in the Cerebellum and essential for motor coordination, have shed some light on the Coding Problem. Those experimental results have illustrated that the memory of time interval duration can be encoded within individual Purkinje cells, and does not require a whole neuronal population [12, 13]. In addition, the stored time memory can be accessed and changed anytime. This result has also challenged the prevailing doctrine of Hebbian learning by showing that traditional changes of synaptic strength alone cannot explain the Purkinje cell response after learning [14].

Purkinje cells can learn to encode a specific time memory through Classical or Pavlovian conditioning. This kind of Associative learning can occur when a biologically potent stimulus, such as food, is paired with a neutral stimulus, such as a bell, that precedes it. Depending upon the response the potent stimulus elicits, e.g., saliva flow, and the exact protocol followed, Classical Conditioning can be categorized into various kinds. One of them being classical motor conditioning, such as the eye blink conditioning, where a neutral conditional stimulus (CS) in the form of a light or a sound can trigger an eye blink response before the onset of an unconditional stimulus (US) that elicits a blink reflex response [15, 16]. In other words, CS triggers a response that predicts the time of arrival of the US. Such a conditional response appears after successive training sessions, where a CS is followed by an US after a fixed time interval “T”, called the interstimulus time interval (ISI) [17]. At the cellular level, the eye blink response is causally related to a suppression of the tonic firing of individual Purkinje cells, which regulate the activity of ocular muscles [17, 18]. Because of such causal connection, the suppression of the firing rate of the Purkinje cell is termed as the conditional response of the Purkinje cell.

Previous mechanistic explanations considered Long-term Depression (LTD) of selective synapses between parallel fibers and Purkinje cells (pf-PC) as the main mechanism behind the conditional response in the Purkinje cell [19]. Based on the widely accepted Marr-Albus model of the cerebellum [20, 21], this suggests that the time memory of the response is encoded within the network dynamics of Granule cell neurons and inhibitory interneurons, found within the molecular layer of the Cerebellum between Mossy fibers and Purkinje cells. However, recent experiments on ferrets were able to identify the source of the conditional response at the level of *individual* Purkinje cells by showing that the direct stimulation of parallel fibers and climbing fibers using electrodes was sufficient for Purkinje cells to learn the specific time interval duration [12]. These experiments also showed that a glutamate-based metabotropic receptor type 7 (mGluR_7_) which resides on Purkinje cells synapses, initiates the conditional response [13] by opening G-protein coupled Inward Rectifier Potassium (GIRK) ion channels [22]. This implies that there exists a specific biochemical mechanism within the Purkinje cell that can encode and store temporal information.

Unlike other memory formation mechanisms requiring neuronal assemblies, temporal signatures can be encoded within a single Purkinje cell, but the specific mechanism remains poorly understood. Here, we propose a biochemical description, based on past experimental findings, that is able to explain time memory formation, consolidation and access.

### Biochemical description

As mentioned above, the conditional response at the level of an individual Purkinje cell appears after several repetitions of two stimuli: A CS from the parallel fibers followed by an US from the climbing fiber after a fixed ISI. A sufficient condition for the learning process to be called completed is that a CS without an applied US can initiate the conditional response — the suppression of the tonic firing rate — with the given ISI. Below we discuss the conditional response of the Purkinje cell and the associated detailed biochemistry we propose. We separate it into three parts: during training, after training and training with different ISIs. The first part focuses on two questions: What makes a Purkinje cell learn a conditional response, and how does the cell learn a conditional response of a specific duration? The later two parts describe the most crucial aspects of the conditional response, i.e., its formation after training, along with other features of the conditional response, which were experimentally observed.

#### During training

What makes a Purkinje cell learn a conditional response? The activation of the conditional response was found to be initiated by the activation of mGluR_7_ receptors [13]. Purkinje cells express mGluR_7_ receptors on their entire cell body and dendritic branches [23], yet no conditional response was observed before training [12]. The fact that in the presence of CS Purkinje cells behave similarly before training and after training if mGluR_7_ antagonists such as 6-(4-Methoxyphenyl)-5-methyl-3-(4-pyridinyl)-isoxazolo[4,5-c]pyridin-4(5H)-one hydrochloride (MMPIP) or LY341495 are present implies that the learning of the conditional response is associated with the expression of mGluR_7_ receptors on the synapse. In other words, during training mGluR_7_ receptors are being transported from the perisynaptic zone to the postsynaptic zone of the synapse via some biochemical mechanism that is activated during training. Once placed on the synapse, mGluR_7_ receptors activate G_*i/o*_ type G-proteins in the presence of glutamate from the parallel fibers. In turn, the G_*βγ*_ subunits of the G_*i/o*_ type G-proteins activate the GIRK ion channels [22, 24]. However, as alternative hypotheses to the absence of mGluR_7_ from the postsynaptic zone before training two other options are conceivable: 1) GIRK ion channels are absent at the synapse; or 2) the expression of G_*i/o*_ type G-proteins was low at the synapse. However, we can rule these out. Immunohistochemistry analysis showed the presence of GIRK subtypes GIRK2/3 ion channels on PC synapses — which are innervated by parallel fibers — in the absence of any kind of prior conditional training [25]. This rules out 1). If 2) were true and the G-protein expression would change during training, this would affect not only the conditional response profile but also various other physiological properties of the Purkinje cell. This is because different types of G-proteins play crucial roles in signal transductions and provide various physiological properties to the cell [26]. Since no change in the tonic firing rate has been observed before and after conditional training [12], we believe that other physiological properties of the cell may also remain unaltered. Thus, the translocation of mGluR_7_ receptors to the synapse is the most likely result of the training and we assume that the amount of other proteins such as GIRK ion channels and G-protein is constant for all different durations of conditional training.

How does the Purkinje cell learn a conditional response of a specific duration? To train a Purkinje cell to memorize a specific duration “T”, two stimuli separated by the same duration, called the Interstimulus Interval (ISI), are required. The first stimulus must come from the parallel fibers and the second stimulus comes from the climbing fiber [17]. Generally, translocation of mGluR type receptors to and from the synapse happens by Clathrin-mediated Endocytosis (CME) [27]. Also, G-protein coupled receptor kinases (GRKs) and in some cases protein kinases such as Protein kinase C (PKC) can phosphorylate receptors and initiate their endocytosis and translocation via CME [27]. Therefore, we propose that the PKC initiates trafficking of mGluR_7_ receptor via CME. This is supported by the fact that the presence of two stimuli, one from the parallel fiber and the other one from the climbing fiber, also makes PKC activation most favorable [28]. In particular, the presence of either one of the two stimuli is not sufficient for the Purkinje cell to learn the conditional response [17]. While dendritic spines of Purkinje cells — which parallel fibers are innervating — express mGluR_1_ receptors [29] and those receptors can potentially activate PKC [30], evidently PKC activation is not the downstream effect of mGluR_1_ receptor activation alone [31, 32]. Similarly, climbing fiber stimuli alone cannot activate PKC. While Long-Term Depression (LTD) of pf-PC synapses occurs via PKC activation [33], it only happens when both parallel fibers and climbing fiber are active [28]. Hence, PKC can become active during training and helps in translocation of mGluR_7_ receptors to the synapse. However, PKC alone is not responsible for the translocation of mGluR_7_ receptors as Purkinje cells cannot be trained for ISI durations shorter than 100ms [34] but PKC can become active even when both CS and US occur at the same time. Currently, we cannot make any suggestion for proteins, which might be involved in addition to PKC during conditional learning.

To ensure storage of a specific time duration memory, such translocation process must stop after some time. This can happen by inhibiting PKC activation, which can be achieved by either removal of mGluR_1_ receptors from the synapse or by preventing the rise of the intracellular Ca^+2^ ion concentration within the synapse. The latter option is the most suitable one because translocated mGluR_7_ receptors at the synapse can open GIRK ion channels, which will drop the membrane potential and thus prevent opening of Voltage-gated Ca^+2^ ion channels. Activation of GIRK ion channels causing a drop in tonic firing rate during training has been observed in experiments [35, 36]. Moreover, if mGluR_1_ receptors were to be removed from the synapse, then the retraining process with a different ISI would not happen as there would be none or very few mGluR_1_ receptors left on the synapse to produce Diacylglycerol (DAG), a necessary membrane-bound biomolecule for PKC activation [30]. As Purkinje cells can be retrained [36], the amount of mGluR_1_ receptors cannot decrease on the synapse. Therefore, we conclude that as training progresses the intracellular Ca+^2^ ion concentration decreases to a level that is no longer sufficient to activate PKC, which prevents further translocation of mGluR_7_ receptors to the synapse and so a steady state will be reached. When a steady state has been reached, then we can say that the Purkinje cell has learned the conditional response of duration “T” as shown in fig.(1). This mechanism of learning also suggests that the training period will be longer for a long duration conditional response as observed in the experiment [12]. As the net amount of the receptor translocated during training depends upon its net duration, long duration training means more transportation of the receptor to the synapse, which produces a long duration conditional response. We will explain how a higher amount of receptors can produce a long duration conditional response below.

**Fig 1.**
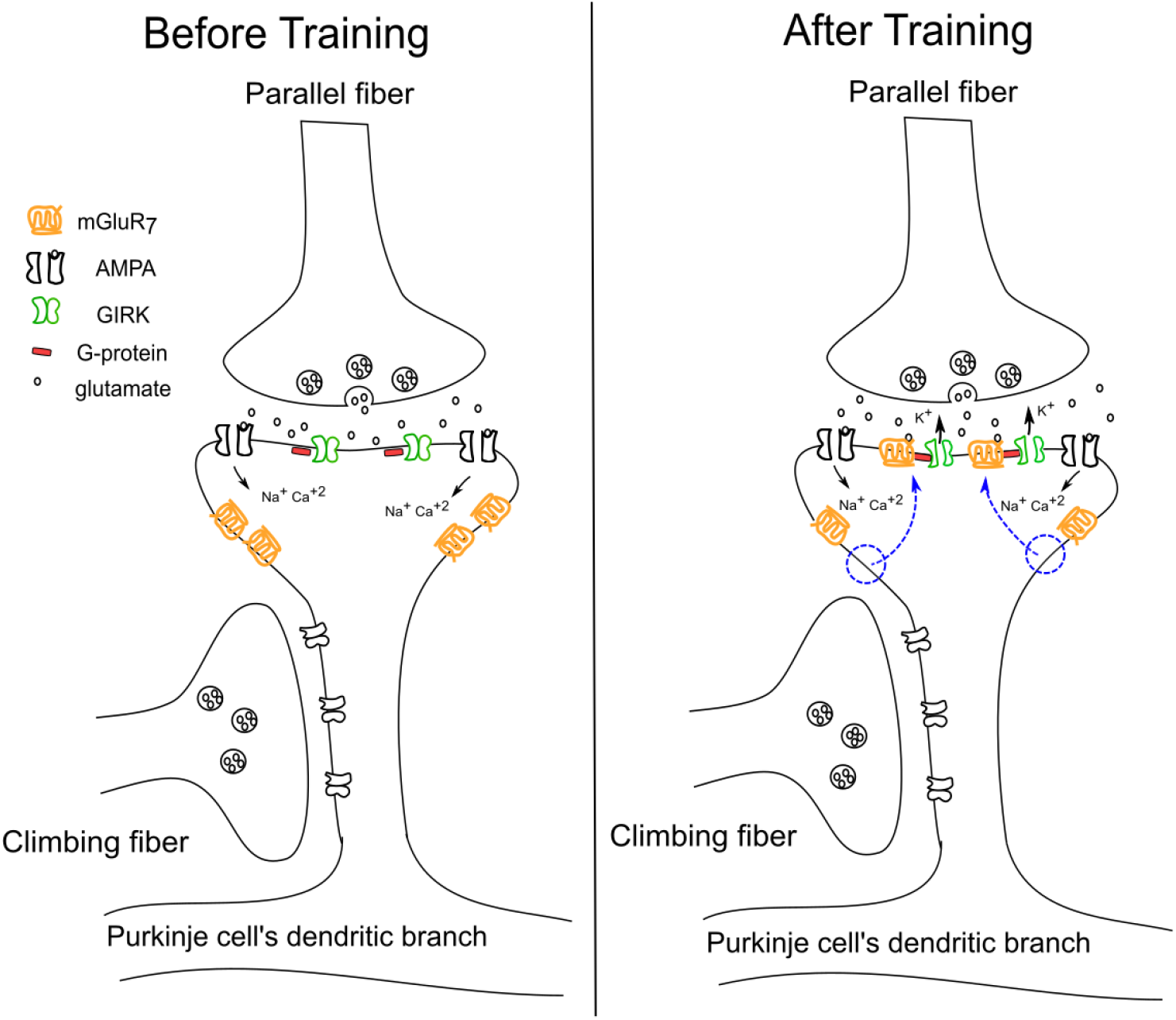
mGluR_7_ receptor distribution before and after conditional training in the Purkinje cell. Before training, mGluR_7_ receptors are localised at perisynaptic areas of the synapses. After training, as pointed out by the blue arrows, these receptors localised themselves at the postsynaptic area of the synapse via CME.

#### After training

The conditional response with a duration of hundreds of milliseconds can be initiated by a CS of as little as 20 milliseconds duration [12]. This means that just the activation of the mGluR_7_ receptors by CS is enough to initiate the conditional response. In addition, the dynamics of the G-protein activation by mGluR_7_ receptors and the binding of G-protein subunits to the GIRK ion channels are usually too slow to explain the fast dynamics of the conditional response initiation observed in the experiment [12]. Such fast activation of GIRK ion channels correlates with some past studies. It was proposed and later verified experimentally that the mGluR_7_ receptor forms a protein complex with the G-protein of G_*i/o*_ type, which is located in direct vicinity to a GIRK ion channel as facilitated by a Regulator of G-protein signaling protein 8 (RGS8) [37, 38]. RGS8 proteins are expressed in dendritic spines of the Purkinje cell [39] and they have a special property to accelerate both activation and deactivation of the G-protein causing fast opening and closing of GIRK ion channels [38]. In particular, it has two domains.

RGS8’s core domain is responsible for rapid deactivation of the G-protein. An additional N-terminal domain helps in the rapid G-protein activation, possibly by enhancing the coupling between G-protein and the receptor [40]. Thus, the fast dynamics of the conditional response is possible by forming a protein complex of mGluR_7_ receptor, G-protein and GIRK ion channel facilitated by a RGS8 protein.

There are two additional important properties of the conditional response, which we must consider: 1) The conditional response is lost after repetitive CS [12], and 2) the conditional response is independent of CS duration. These two properties are in fact related to each other. The former implies the removal of mGluR_7_ receptors from the synapse and the latter implies that some protein blocks the receptor’s active site to prevent reactivation of the conditional response. Dephosphorylation of the mGluR_7_ receptor by Protein Phosphatase 1 (PP1), which causes their rapid internalization [41], can explain both these properties of the conditional response. Rapid internalization of any receptor is initiated by the binding of a protein called Arrestin protein, which prevents the receptor to transmit any signal further [30]. Since CS activates the conditional response, it implies that PP1 is inactive before the conditional response. Furthermore, the conditional response is mediated via activation of the G_*i/o*_ type G-protein whose G*_α_* subunit blocks the production of Cyclic adenosine monophosphate (cAMP) by Acetyl Cyclase (AC) and, thus, results in reduced Protein Kinase A (PKA) activity. Also, due to the tonic firing of the Purkinje cell, Calmodulin, which regulates intracellular Ca^+2^ ion concentration [42], can stimulate Acetyl cyclase (AC) [43] to produce cAMP molecules and increase PKA activity. It is also known that PKA can phosphorylate mGluR_7_ receptors [44] as well as PP1 regulatory proteins such as Dopamine- and cAMP-Regulated neuronal Phosphoprotein (DARPP-32) or Inhibitor-1 (I-1) [45], which prevents dephosphorylation of receptors. Thus, phosphorylation of the receptor and inhibition of PP1 activity by PKA help in the retention of the memory for a long time. As PKA is essential for the conditional response, this protein could bind to the receptor via a special PKA anchoring protein called A-kinase anchoring proteins (AKAP) [46]. Similarly, PP1 can also bind close to the receptor via another scaffold protein namely Spinophilin [47], which is expressed in dendritic spines of neurons across various regions of the brain including Cerebellum [48]. Such a close agglomeration of various proteins together forms a protein complex, which is self-sufficient in its own regulation and dynamics.

In short, the underlying biochemical mechanims of the conditional response can be described as follows. Before CS, the PP1 protein is inactive because of PKA activity. The release of glutamate during CS activates mGluR_7_ receptors on the Purkinje cell synapses [step 1 of Fig. (2)], which in turn activates G-proteins [step 2 of Fig. (2)]. Each unit of G-protein splits into a G*_α_* subunit and a G_*βγ*_ subunit. One unit of G*_α_* subunit binds to an AC enzyme to block the production of cAMP molecules. This in turn deactivates PKA as Phosphodiesterase enzyme (PDE) hydrolyses the remaining cAMP molecules [30] [step 3 of Fig. (2)]. At the same time the G_*βγ*_ subunit binds to the GIRK ion channel, which becomes fully active upon binding of four G_*βγ*_ subunits [24]. As PKA activity decreases, PP1 activity rises due to dephosphorylation of DARPP-32 or I-1 by Protein Phosphatases such as PP2A [45, 49] [step 4 of Fig. (2)]. The rise of PP1 activity causes dephosphorylation of mGluR_7_ receptors [step 5 of Fig. (2)] and initiates their rapid internalization. However, rapid internalization of a receptor is still a slow process compared to the conditional response as it involves many protein interactions and, hence, the receptor is not immediately displaced from the synapse after dephosphorylation. However, after dephosphorylation, Arrestin protein blocks the active site of the mGluR_7_ receptor to prevent reactivation of the G-protein [50]. After receptor dephosphorylation, the active G-protein is deactivated by the RGS8 protein [step 6 of Fig. (2)]. As G-protein activity reduces, GIRK ion channels also shut down. In the absence of active G-protein, PKA activity begins to rise again [step 7 of Fig. (2)] due to rise in activity of AC enzymes in the presence of Calmodulin. Active PKA deactivates PP1 [step 8 of Fig. (2)] by phosphorylating DARPP-32 or I-1 and finally, PKA also phosphorylates mGluR_7_ receptors [step 9 of Fig. (2)] to prevent their internalization and prepare the Purkinje cell for another conditional response. It is likely that the reactivation of PKA takes some time, which might explain why CS cannot initiate another conditional response while CS is still on.

**Fig 2.**
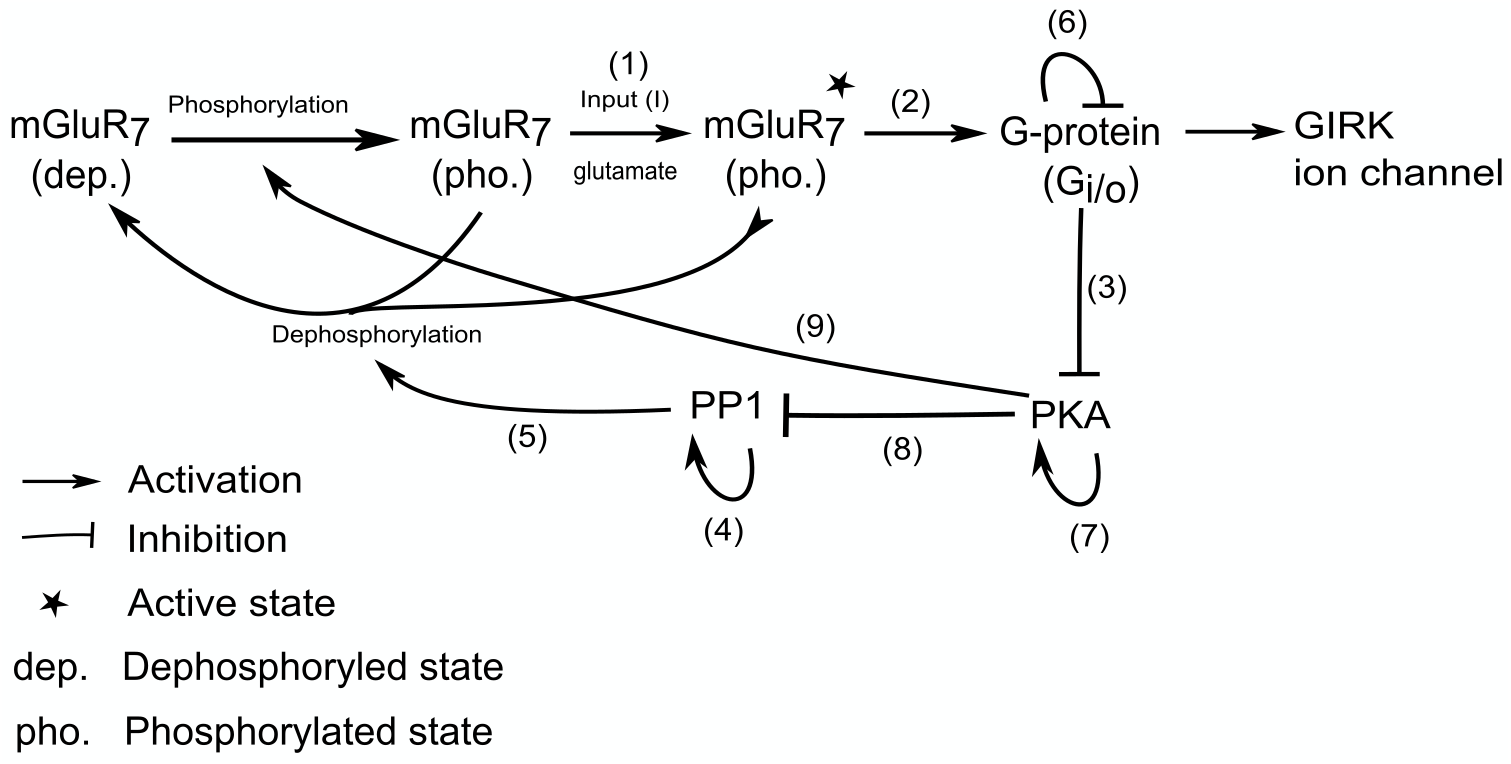
Interactions between different biochemicals involved in our proposed mechanism. The numbers on the top of the arrows highlight the order in which the different reactions occur during the conditional response. Conditional response initiates with the release of glutamate from parallel fibers denoted by I as input in (1), which activates mGluR_7_ receptors. In (2), active receptors activate G-proteins, which deactivate PKA through (3). As PKA activity reduces, PP1 activity rises through (4) causing dephosphorylation of the receptor (5). As receptor activity reduces, RGS8 reduces G-protein activity (6), which allows PKA activity to rise again (7). Active PKA will deactivate PP1 (8) and lastly phosphorylate dephosphorylated receptors to prevent their rapid internalization (9).

In fig.(2), the rate at which GIRK ion channels open and close depends upon the rate at which intermediate reactions occur. In other words, the time memory of the training is stored within the effective rate constants arising from these reactions. In a complete cycle of GIRK ion channel activation and deactivation, altering only effective rate constants for both activation and deactivation processes is sufficient to store a different time memory of the conditional response.

#### Training with different ISI duration and time-encoding protein complexes

Training with a different ISI duration means storage of a different time memory. As discussed above, the time memory is encoded within effective rate constants of the biochemical reactions, which regulate the gating dynamics of the GIRK ion channel and altering these rate constants means storing a different time memory. However, there are two additional questions we need to answer in order to get a complete understanding of time memory storage in biochemical reactions: 1) How do these biochemical reactions get tuned so finely to store a specific time duration memory? 2) Among all possible effective rate constants of the proposed biochemical mechanism, which rate constants are most likely to get affected by choosing a different ISI for the training?

The reason behind 1) is that there are several GIRK ion channels present at the synapse. Each GIRK ion channel requires four units of G_*βγ*_ subunits to open completely [24]. This means that each GIRK ion channel forms a protein complex with four units of G-protein, receptor and RGS8 protein along with PKA and PP1 proteins with their anchoring proteins together. As each of these protein complexes has their own intrinsic dynamics, which regulate how fast the GIRK ion channel opens and closes upon stimulation, we can call each of these protein complexes “Time-Encoding protein Complexes” (TEC). Within each TEC, the rate of G-protein activation by the receptor and the rate of binding of G-protein subunits to the GIRK ion channel decide the overall rate of opening of GIRK ion channels i.e., the onset of conditional response. After the onset of the conditional response the rates of PKA deactivation, PP1 activation, dephosphorylation of the receptor and the deactivation of G-protein by RGS8 decide the overall duration of the conditional response since at the end of these biochemical reactions the GIRK ion channel begins to close. Thus, each TEC encodes time information of the conditional response completely in terms of effective rate constants of different biochemical interactions and stores this time memory by forming a protein complex. Formation of a protein complex as TEC ensures strong consolidation of memory with less chances of errors in the information storage. If the rates were to be changed so would the memory as well. The rates can be affected by the translocation of extra mGluR_7_ receptors to the synapse during conditional training. These extra mGluR_7_ receptors can form clusters with receptors — which are part of TEC — with the help of a scaffold protein, Protein Interacting with C Kinase - 1 (PICK1) [51]. Such cluster formation can affect TEC’s intrinsic dynamical properties by influencing the protein interaction of the mGluR_7_ receptor with the G-protein facilitated by RGS8. As a result, RGS8’s ability to accelerate the dynamics of the conditional response might be affected, which results in a delayed onset of the conditional response. Such clustering of receptors can also affect the dynamics of the PKA protein anchored close to the receptor via the AKAP protein, thus affecting also the time duration of the conditional response. To summarize, we propose that at individual synapses the interaction of extra mGluR_7_ receptors with TECs can affect the dynamics of TECs and collectively these varied TEC units help to produce the conditional response of any specific time duration in the Purkinje cell.

Interestingly, time memories stored at synapses in the form of TECs are not permanent but can be altered or edited through retraining. Retraining can happen via two ways, (i) erase the memory first and then store another memory by training with a different ISI interval, or (ii) retrain the Purkinje cell with a different ISI without erasing the previous memory. Experimentally, time memory can be erased by repeating unpaired representation of CS and US several times [12], which can also be explained by our proposed mechanism. Due to the action of PP1 on the receptor, every time CS initiates a conditional response, some of the receptors at the synapse might undergo rapid internalization from which phosphorylation of the receptor by PKA cannot bring them back and, hence, these receptors will be removed from the synapse. Because of rapid internalization, retraining with the same or a different ISI will be faster as many receptors are close to the synapse. This rapid relearning phenomenon is called “Saving” and it takes only a few minutes to recall the old memory [17]. If retraining with a different ISI is performed without erasing the old memory then the conditional response profile differs in terms of the onset of the conditional response while the duration remains intact when compared with the case where retraining does involve erasing the old memory prior to retraining [17, 52]. This difference implies that there might be some membrane-bound proteins or a scaffold protein such as PICK1 that are interacting with TECs. If the two ISI durations differ significantly then surprisingly, the Purkinje cell pauses twice in the presence of a sufficiently long duration CS [12]. Depending upon the time difference between the two different ISIs, different conditional responses were observed implying that there might be additional interactions among TECs, which give rise to a wide variety of conditional responses [53].

From the above description, it follows that in principle a single synapse can completely contain a time duration memory, which can be altered through retraining. However, a single synapse probably will not be sufficient to suppress the tonic firing rate of the whole Purkinje cell. This is because the spontaneous tonic firing rate of the Purkinje cell [54] appears to be due to voltage-dependent resurgent Na^+^ ion channels, which are distributed over the *entire* somata and dendritic regions of the cell [55, 56]. The resurgent Na^+^ channels have the property to become active and inactive during depolarization, as well as to reactivate during repolarization due to the presence of a positive membrane potential. The latter results in a spontaneous rapid sequence of action potentials [57]. In order to suppress the spontaneous firing rate of the cell, these resurgent Na^+^ ion channels need to be deactivated by the hyperpolarizing membrane potential of the entire cell. Activation of GIRK ion channels by CS can hyperpolarize the membrane at a synaptic regions and deactivate resurgent Na^+^ ion channels near those synaptic region. Their deactivation will lower the overall membrane potential and the activation of multiple pf-PC synapses distributed over the dendrites can overcome the resurgent Na^+^ current and suppresses tonic firing rate of the cell. Thus, a finite fraction of the total pf-PC synapses can produce a suppression in tonic firing rate of the Purkinje cell for a specific duration and the corresponding memory is encoded at the respective synapses.

In the next section, we will introduce and discuss a mathematical model of our proposed biochemical mechanism and subsequently make few predictions that can be tested experimentally.

### Mathematical model

To model the conditional response behavior of the Purkinje cell, we start with an established dynamical model of the Purkinje cell which incorporates many properties of the cell within a realistic biophysical framework [58], see Materials and Methods for details. Before training, GIRK ion channels cannot be opened because mGluR_7_ receptors are not present at the synapse. However, after training, mGluR_7_ receptors are present at the synapse to open GIRK ion channels. Therefore, adding an additional term for the gating of the GIRK ion channel in the Purkinje cell model will allow it to exhibit the conditional response. As GIRK ion channels reside at synapses, the additional term of GIRK ion channel gating must be added in the dendritic equation of the Purkinje cell model. Eq.(1) defines the gating of the GIRK ion channels, in which *g_GIRK_* is the net conductance of GIRK ion channels per unit area, *h_GIRK_* is the gating parameter and *V_GIRK_* is the voltage dependence of the GIRK ion channel obtained from the I-V characterstics curve [59]. Since no experimentally measured value exist for *g_GIRK_* to the best of our knowledge, we choose its value to match the conditional response observed experimentally.

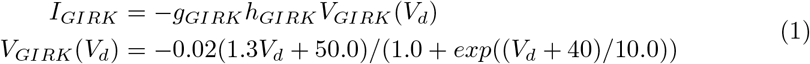

Gating of the GIRK ion channel depends upon the availability of the Phosphatidylinositol 4,5-bisphosphate (PIP_2_) molecules [24]. This molecule has low affinity for the GIRK ion channel but binds efficiently after binding of G_*βγ*_ subunit to the GIRK ion channel. The amount of PIP_2_ on the synaptic membrane is low but its quantity is often replenished by various biochemical processes to maintain its concentration fairly constant upon consumption or degradation [60]. Therefore, the amount of active G_*βγ*_ subunits can determine the gating dynamics of the GIRK ion channel. As G-protein is closely associated with the GIRK ion channel, we can assume fast binding of the G_*βγ*_ subunit to the GIRK ion channel. With these assumptions, we can equate the normalized G-protein activity with the GIRK ion channel gating parameter h_GIRK_ as shown in eq.(6) below.

G-protein activity depends upon the activity of the mGluR_7_ receptor along with other proteins as shown in fig.(2), which self-orgainze to form discrete units of TECs. Since we don’t know the number of TECs and their detailed intrinsic dynamics, we choose to model the collective dynamics of TECs and different biochemical interactions within them in an effective way. Hence, instead of using discrete variables for the activity of different biochemicals, we use continuous variables to capture the “average” dynamics of different biochemicals by considering all TECs together. In addition, AKAP proteins — which anchor PKA close to cAMP production machineries — do accelerate the activation of PKA, but not sufficiently fast to generate a conditional response [61, 62]. It is possible that other proteins such as Homer Proteins might also be involved in the TEC, facilitating cross-talk between various target proteins [63]. Thus, due to lack of knowledge of various protein interactions and their strengths within individual TECs, we only attempt to create a conceptual minimal model whose aim is to reproduce features of the conditional response, namely 1) the conditional response should be independent of CS duration, and 2) changing the dynamics of PKA and G-protein should be sufficient to produce conditional responses of different durations. Our conceptual minimal model consists of four main biochemicals - mGluR_7_, G-protein, PKA and PP1. In order to simplify and minimize the number of parameters, we used nonlinear terms to model their overall behaviour as observed *in vivo*.

The nondimensional dynamical equations for the proposed biochemical mechanism within individual TECs are as follows:

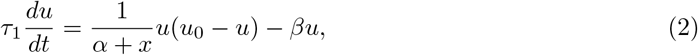

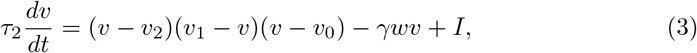

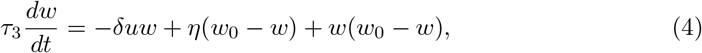

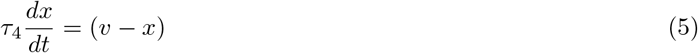

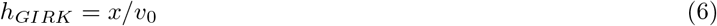

where *u, v, w* and *x* are the activities of PKA, mGluR_7_ receptor, PP1 and G-protein respectively. In the above model, all the parameters and variables are positive and dimensionless quantities except for *τ_i_* for *i* = 1, 2, 3, 4, which have dimension of time.

Eqs.(2 – 5) match the pictorial diagram shown in Fig. 3, which depicts various variables and their dependencies. In order to understand various terms within each equation, let us focus on each equation individually. Before CS, when G-protein is still inactive, i.e., *x* ~ 0, AC enzyme produces cAMP molecules facilitated by the Calmodulin protein. As the activity of AC enzyme increases, cAMP production also increases, which increases PKA activity. This behaviour of PKA activity is modeled in eq.(2) by the first term *u*(*u*_0_ − *u*)/*α* for *x* = 0, where *u*_0_ is the maximum PKA activity possible. Active PKA phosphorylates phosphodiesterase enzyme (PDE), which hydrolyses cAMP molecules to Adenosine monophosphate (AMP) [64]. Activity of PDE depends upon the activity of PKA since it can be dephosphorylated by Protein Phosphatases such as PP2A. Thus, PDE activity depends on the PKA activity to reduce the net PKA activity. This behaviour of PDE is modeled by the second term −*βu* in eq.(2), where the parameter *β* signifies the strength of the PDE action on PKA deactivation. Upon parallel fiber stimulation, glutamate activates the mGluR_7_ receptor, which activates G-protein to produce a G_α_ subunit to block the cAMP molecule production. This behaviour is modeled by the prefactor 1/(*α* + *x*) of *u*(*u*_0_ − *u*) in eq.(2), where *x* denotes G-protein activity. The prefactor corresponds to blocking of AC enzyme, which is obtained from Hill’s equation with Hill’s coefficient equal to 1 as only one unit of G*_α_* protein binds to AC. For more details on Hill’s equation, see Materials and Methods. The constant *α* denotes the disassociation constant *K_d_* of AC and G*_α_* subunit and it has a small value due to their strong bonding. *τ*_1_ signifies the overall time scale of the PKA dynamics. Since according to the proposed mechanism different conditional responses are the result of changes in the dynamics of PKA activation and deactivation, the value of *τ*_1_ will increase or decrease for a conditional response of longer or shorter duration, respectively.

**Fig 3.**
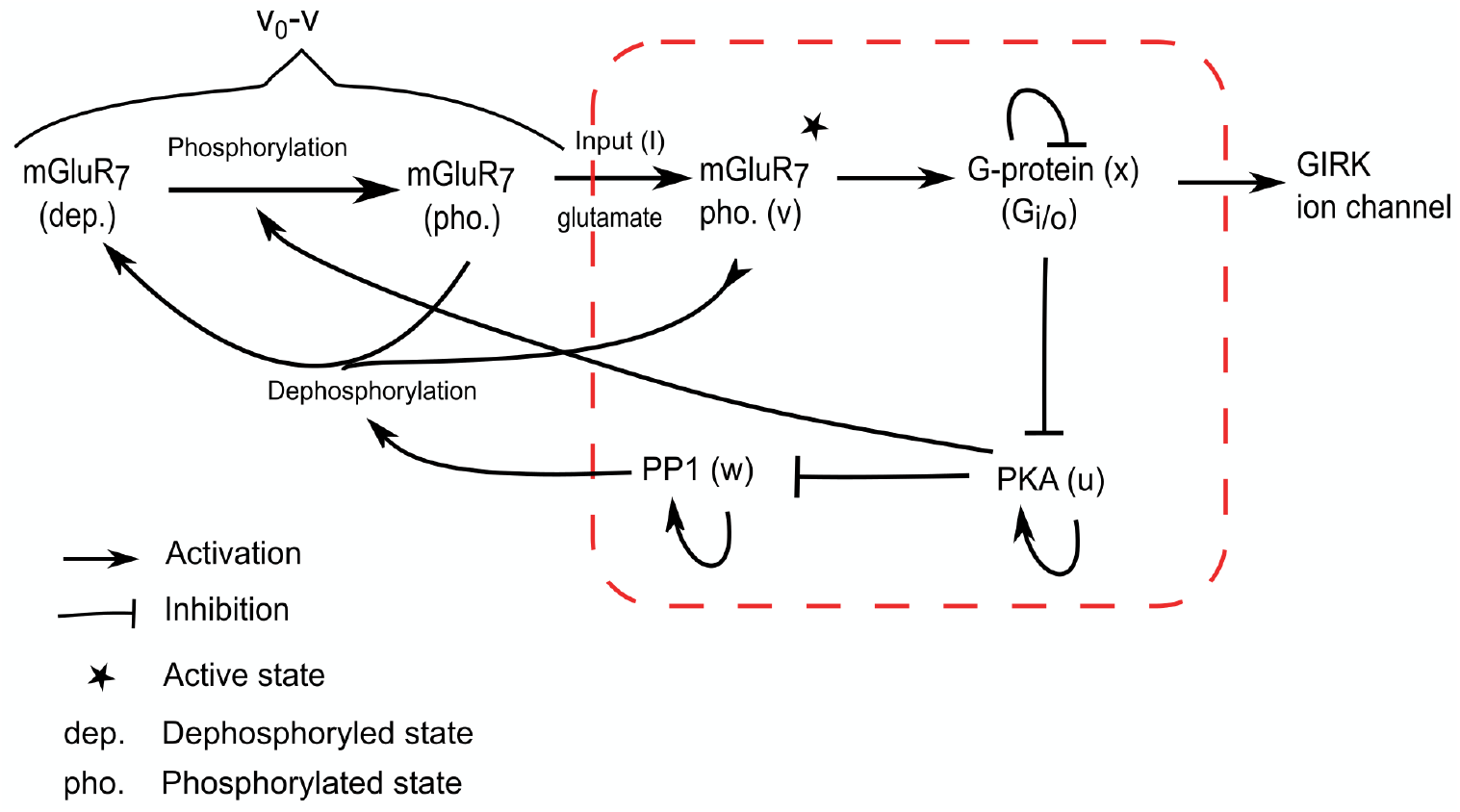
A conceptual minimal model of conditional response in the Purkinje cell.

Eq.(3) models the activity of the mGluR_7_ receptor. The first term is a cubic polynomial, which captures the switching property of the mGluR_7_ receptor corresponding to the unaltered conditional response with changing the CS durations. In the cubic polynomial *v*_0_ ≫ *v*_1_ ≳ *v*_2_, where *v*_0_ signifies the amount of receptors, which are associated with the G-protein, *v*_1_ is the threshold activity that needs to be crossed to initiate the conditional response and *v*_2_ is the net finite intrinsic activity of the receptor [65]. As each mGluR_7_ receptor can form a protein complex with only one unit of G-protein, the value of *v*_0_ remains constant as we assumed that the amount of G-protein is constant for all different conditional response trainings. The values of *v*_1_ and *v*_2_ depend upon intrinsic properties of the receptor itself, which we also assumed to be constant. After deactivation of PKA by G-protein activity, activity of PP1 rises, which dephosphorylates the receptor resulting in the blockade of the receptor’s active site to activate G-proteins further. This is modeled as −*_γwv_* in eq.(3), which denotes the lowering of net receptor activity due to dephosphorylation by PP1. It is a product because PP1 interacts with the mGluR_7_ receptor directly during dephosphorylation. The factor *γ* denotes the strength of the influence of PP1 on mGluR_7_ receptors. As Spinophilin binds PP1 close to the mGluR_7_ receptor by binding to RGS8 [66] and not the receptor, the strength of influence of PP1 on mGluR_7_, i.e., the value of *γ*, can be assumed constant for different conditional responses. Finally, I denotes the strength of the CS in the form of glutamate release from parallel fibers to activate mGluR_7_ receptors and *τ*_2_ signifies the overall time scale of receptor activation and deactivation. The value of *τ*_2_ is taken to be smaller than the fastest onset of the conditional response observed in the experiment [12].

Eq.(4) models the activity of the PP1 protein. Its activity is regulated by PKA as a suppressor by phosphorylating DARPP-32 or I-1 protein, which is modeled as −*δuw*. It is a product because PKA interacts directly with the PP1 regulatory protein, which binds to PP1 to block its activity. The factor *δ* signifies the strength of the PKA influence on PP1 activity. As the activity of PKA decreases, PP1 activity rises by the action of other Phosphatase proteins such as PP2A/B [45] and by itself [67]. The rise due to other Phosphatase proteins is given by *η*(*w*_0_ − *w*), while the rise of PP1 by itself is given by *w*(*w*_0_ − *w*), where *w*_0_ is the maximum activity of PP1 and the factor *η* controls the strength of the influence of other Phosphatase proteins on the rise of PP1 activity. *τ*_3_ signifies the overall time scale of PP1 activation and deactivation. For simplicity, we assume all the variables in this equation to be constant.

Eq.(5) models G-protein activity. Since we assumed that the amount of G-protein on the synapse is constant, net G-protein activity will be the same for all different conditional responses. As a result, G-protein activity will be limited to ‘*v*_0_’, which is the total amount of G-protein present at the synapse and modeled as (*v* − *x*). *τ*_4_ signifies the time scale of activation and deactivation of the G-protein. As different conditional response trainings involve a different amount of mGluR_7_ receptors, the net dynamics of G-protein activation and deactivation by RGS8 is affected by extra mGluR_7_ receptors interacting with TECs. Therefore, depending upon training, the value of *τ*_4_ can be small or big. This will result in a short delay or a long delay in the onset of the conditional response, respectively. When v reduces due to PP1 activity, (*v* − *x*) < 0. This signifies the deactivation of the G-protein due to the action of the RGS8 protein.

In eqs. (3) and (4), the terms −*γwv* and −*δuw* signify the interaction of PP1 with mGluR_7_ and PKA with PP1, respectively. Yet, there are no corresponding terms in eq.(2) of PKA and eq.(4) of PP1 because those interactions are enzymetic in nature and have very short time scales compared to the response, which we are trying to model. Hence, the activity of PKA and PP1 does not change when they interact with other proteins. Note that the *τ_i_* factors on the LHS of each equation make them dimensionless.

## Results

### Properties of the model

Since experimental results have shown that the conditional response is independent of CS durations, the activation of the G-protein must also satisfy this property as it regulates GIRK ion channels. This behavior is indeed captured by our mathematical model. It also successfully captures the dynamics of other biochemicals - PKA, mGluR_7_, PP1 and G-protein as proposed in the mechanism that is shown in Fig. (4). As per our proposed mechanism, before CS, PKA activity is high while activity of mGluR_7_ receptor, G-protein and PP1 is low. Upon CS, mGluR_7_ receptors become active, which in turn activate G-protein. Due to activation of G-protein, PKA activity drops down, which causes a rise in the PP1 activity. When PP1 activity is sufficiently high, it causes deactivation of mGluR_7_ receptors, which then causes deactivation of the G-protein by RGS8. When the stimulus is turned off, activities of various proteins return back to their original states as shown in the top panel of Fig. (4). However, if the stimulus remains on for a long time, then even very low G-protein activity can prevent the rise of PKA activity to a value that would be high enough to block PP1. As a result, PP1 activity will be crucial to cause dephosphorylation of mGluR_7_ receptors and to block their active sites to prevent the initiation of another conditional response as shown in the middle panel of Fig. (4). Very faint G-protein activity in case of long duration CS can be observed in the bottom panel of Fig. (4). This is enough to prevent reactivation of the conditional response.

**Fig 4.**
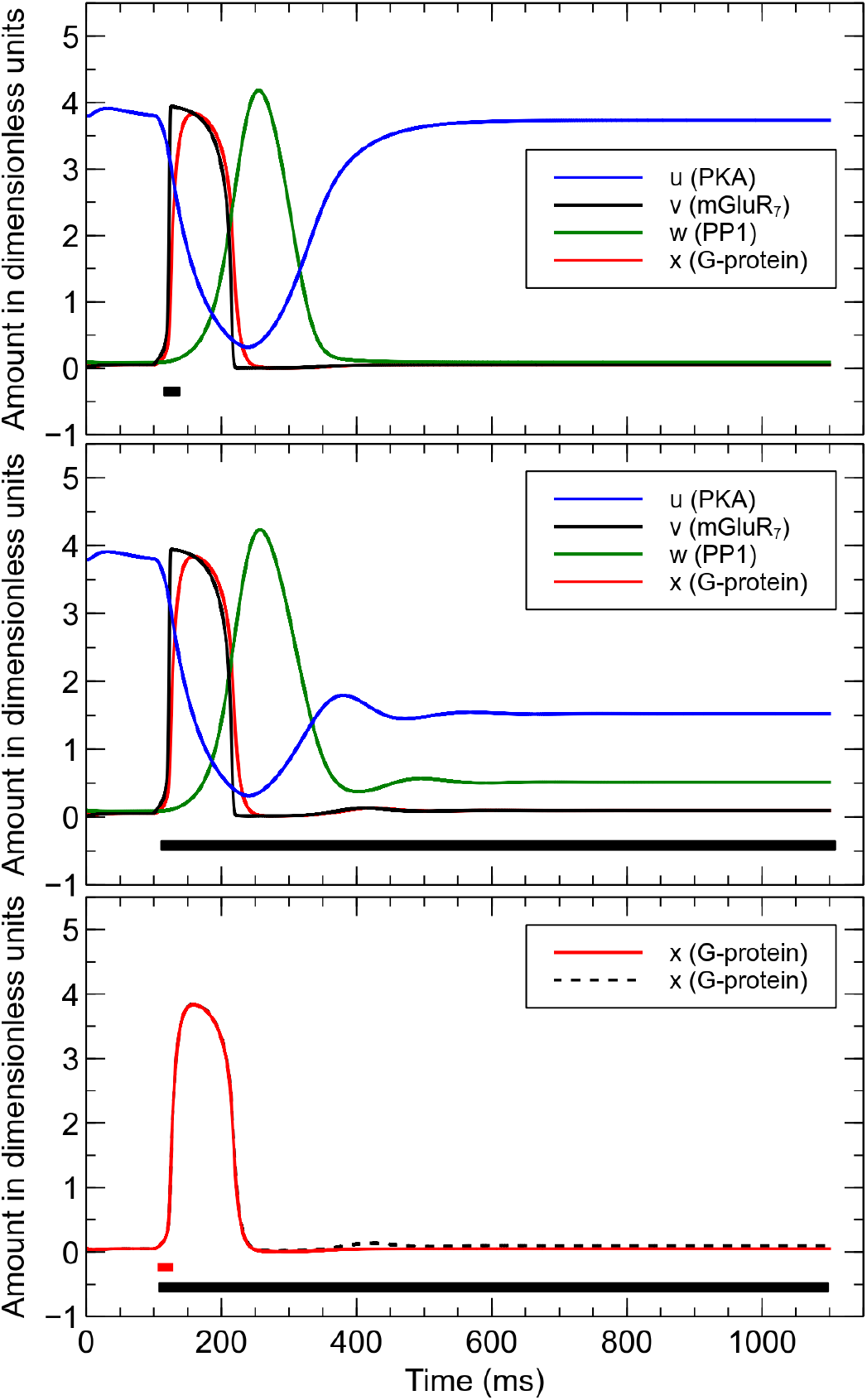
Temporal behaviour of PKA, mGluR_7_, PP1 and G-protein. Time varying dimensionless quantities of PKA, mGluR_7_, PP1 and G-protein upon short (top panel) and long (middle panel) stimulus durations represented by black horizontal bar at the bottom of each panel. In the bottom panel, activity of the G-protein is shown as in the upper two panels but for two different stimulus durations together, indicated by the red and black bars at the bottom. Both responses are almost identical implying that the G-protein activity is indeed independent of stimulus duration. Parameters for our dynamical model of PKA, mGluR_7_, G-protein and PP1 are: *α* = 0.1, *u*_0_ = 11.0, *β* = 48, *v*_0_ = 4.0, *v*_1_ = 1.01, *v*_2_ = 1.0, *γ* = 2.0, *δ* = 5.0, *η* = 0.2, *w*_0_ = 6.0, *I* = 0.1, *τ*_1_ = 2100*ms*, *τ*_2_ = 6*ms*, *τ*_3_ = 60*ms*, *τ*_4_ = 7.9*ms*.

Specifically, the chosen parameter values will determine the specific extreme values of the various variables and their temporal profiles. While these values vary with the chosen parameters, the overall properties of the model will not be affected as long as two features are preserved: i) The G-protein activation remains largely independent of the CS duration, and ii) no oscillatory response emerges. These two features are essential in order to reproduce experimental results. We have verified that these features are preserved over a large range of parameter values for our model. Specifically, changes in size of up to 100% of the values we use in the different figures will render above mentioned two features unchanged. Thus, our conceptual model can robustly capture the main experimental results.

By combining the dynamical equations of mGluR_7_, G-protein, PKA and PP1 with the Purkinje cell model, we can generate the conditional response dynamics of the Purkinje cells as shown in Figs. (5,6). In Fig.(5) we show that the suppression of firing rates during the conditional response of ISI = 150ms is independent of CS durations as observed in the experiments [12]. Result shown in Fig.(5) can be considered as an average response of the firing rate during the conditional response given the deterministic nature of our model.

**Fig 5.**
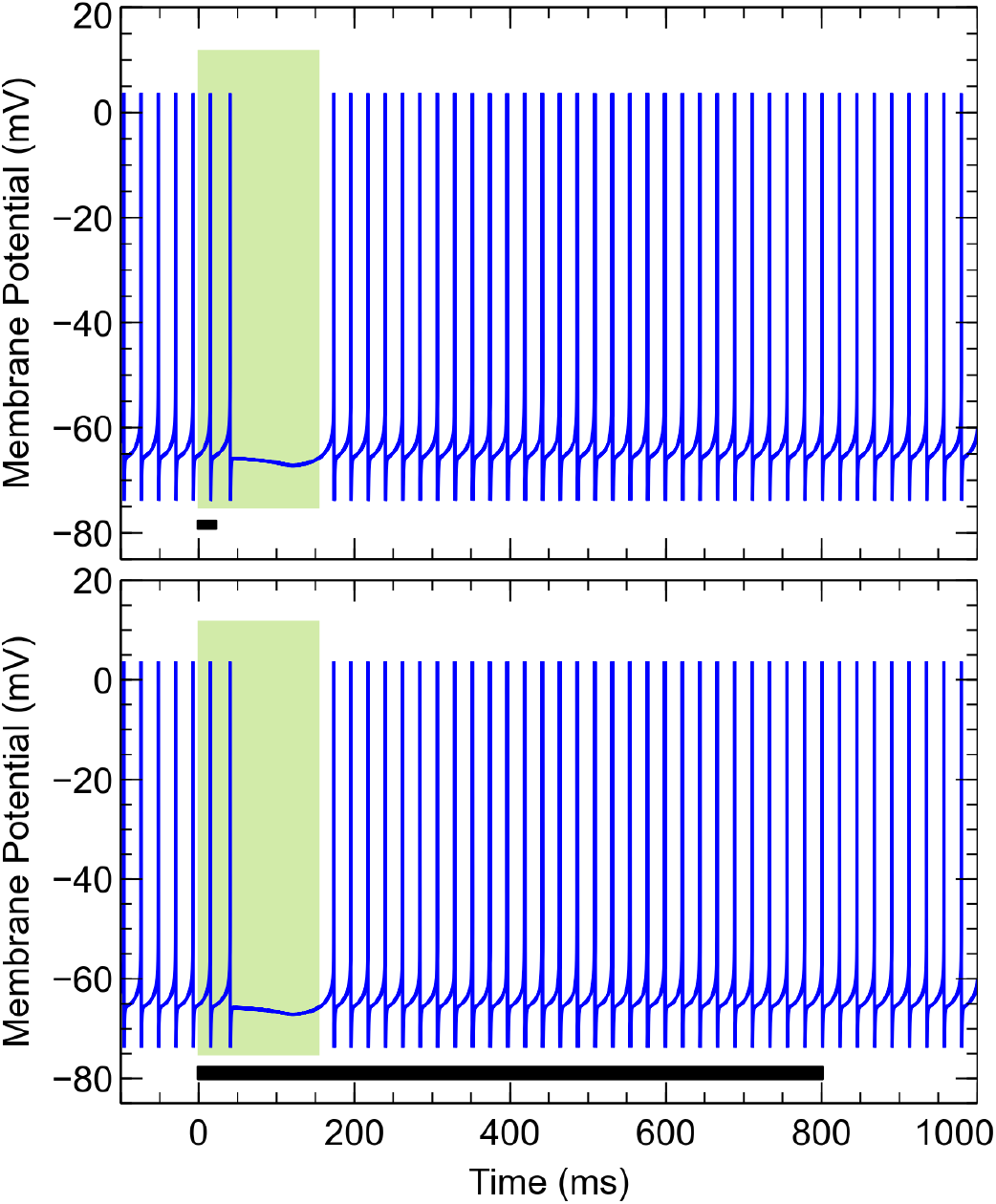
Conditional response in the model is independent of conditional stimulus duration. The width of the light green vertical bar corresponds to the duration of the ISI = 150 msecs and the black bar at the bottom signifies the conditional stimulus duration, which is short in the top panel and long in the bottom panel. Parameters for the dynamical model of PKA, mGluR_7_, G-protein and PP1 are the same as in Fig. 4. For all other parameters of the Purkinje cell model, see Materials and Methods.

In order to obtain a conditional response of longer duration, more mGluR_7_ receptors need to be inserted into the synapse. These extra receptors cause a rise in the value of *τ*_1_ and *τ*_4_ as discussed earlier. Different *τ*_1_ and *τ*_4_ values, which we have used for reproducing different conditional responses, are summarized in Table 1. Fig.(6) shows different long duration conditional responses, which match with the experimental results [12]. Fig.(6) shows the drop in firing rate for ISI = 200ms, while Fig.(6) shows the drop in firing rate for ISI = 400ms as an additional case. Indeed, the three conditional response firing patterns obtained from our model shown in the left panel of Fig. (7) match with the experimental results [13]. In addition, our proposed mechanism also explains why the time-memory remains unaffected in the presence of mGluR_7_ antagonist 6-(4-Methoxyphenyl)-5-methyl-3-(4-pyridinyl)isoxazolo[4,5-c]pyridin-4(5H)- one hydrochloride (MMPIP) as observed in the experiment [13]. Specifically, because of the presence of MMPIP, fewer mGluR_7_ receptors are left to activate GIRK ion channels, which leads to a smaller drop in firing rate. However, reducing the net amount of active mGluR_7_ does not inhibit internal interactions between receptor and other proteins involved in our proposed mechanism. Hence, the time-memory, which is encoded within effective rate constants of biochemical reactions, is unaffected by MMPIP as shown in Fig. (7). Note that in Fig. (7) the action of an increasing dose of MMPIP is simulated by decreasing the value of the parameter *g_GIRK_*. As the corresponding values of *g_GIRK_* have not been measured experimentally as mentioned above, we choose suitable values of gGIRK for the given parameter value of the Purkinje cell model Materials and Methods.

**Table 1.** Model parameters for different conditional responses of the Purkinje cell.

| ISI (ms) | $\tau_1$ (s) | $\tau_4$ (ms) |
| --- | --- | --- |
| 150.0 | 2.1 | 7.9 |
| 200.0 | 7.2 | 33.3 |
| 300.0 | 15.0 | 79.0 |
| 400.0 | 18.0 | 120.0 |

**Fig 6.**
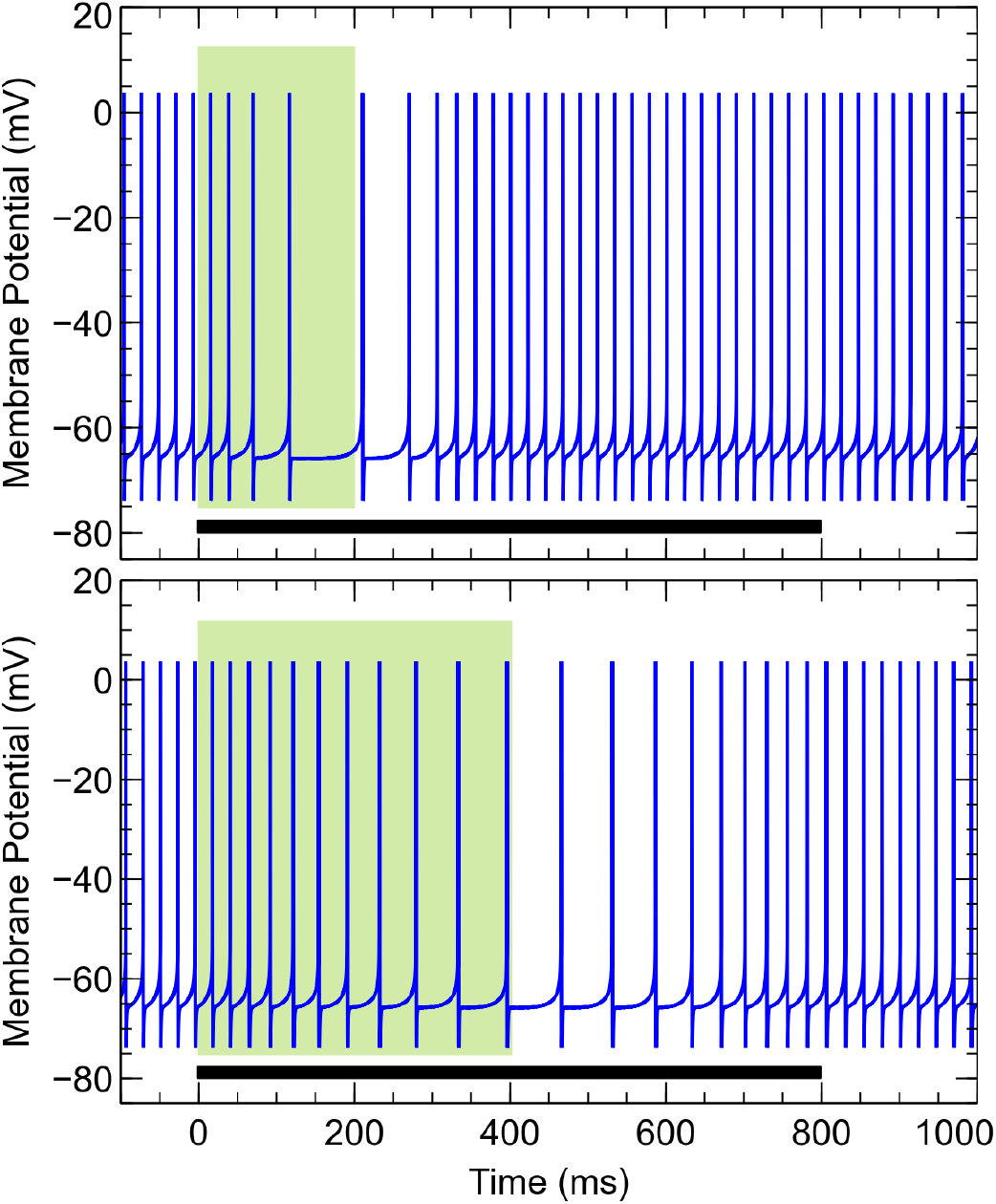
Different conditional responses of the Purkinje cell obtained from the mathematical model. For ISI = 200ms (top panel), the firing rate drops and then rises slowly, which is consistent with the experimental results. The values of *τ*_1_ and *τ*_4_ are 7.2s and 33.3ms, respectively. For higher ISI = 400ms (bottom panel), the drop and rise of the firing rate is observed to be even slower compared to ISI = 200ms at the top panel. The values of *τ*_1_ and *τ*_4_ for ISI = 400ms are 18.0s and 120.0ms, respectively. The width of the light green vertical bar corresponds to the duration of the ISI interval. The black horizontal bar at the bottom represents the conditional stimulus duration.

**Fig 7.**
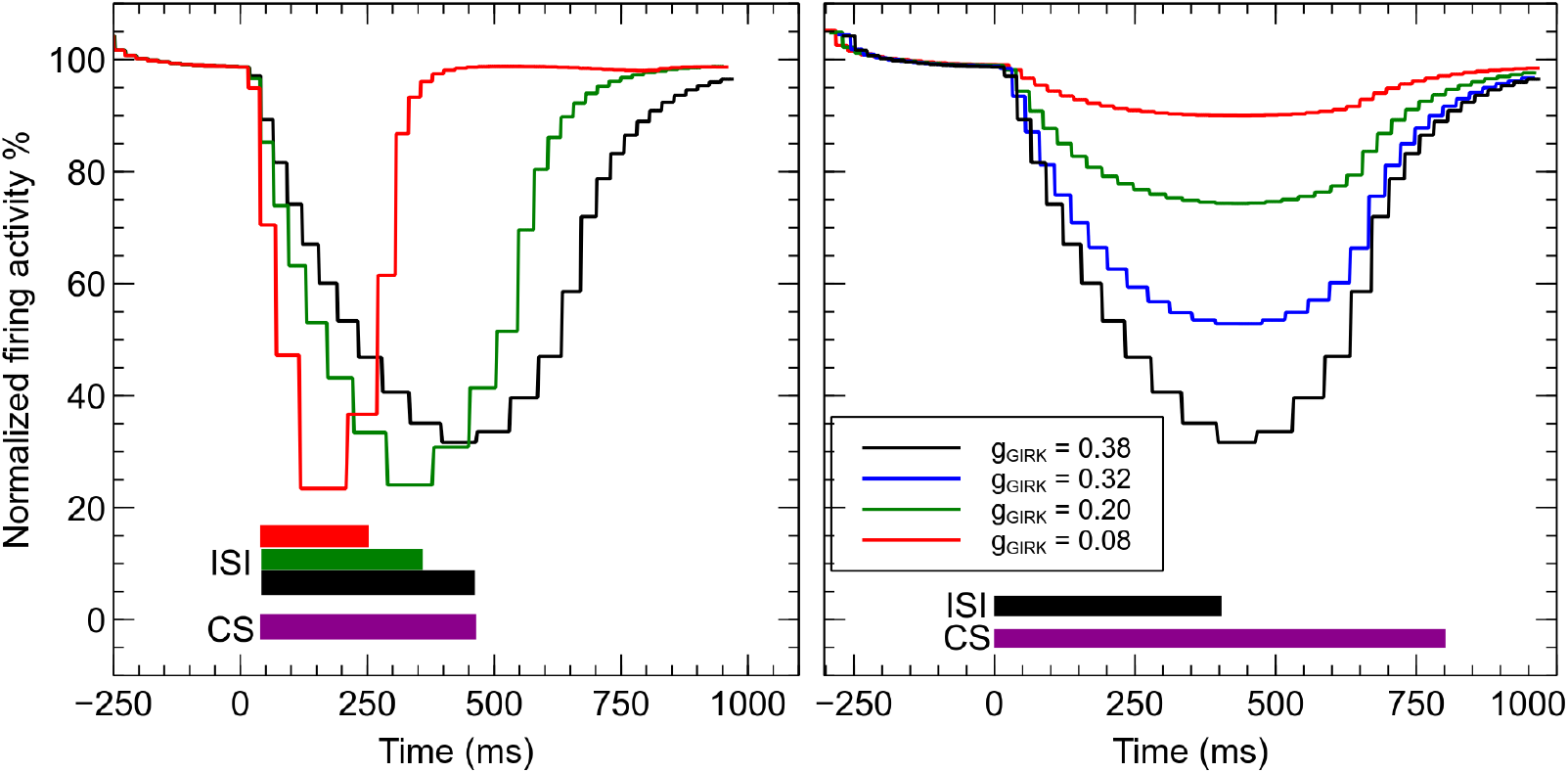
Conditional response profiles for different ISIs and different amounts of MMPIP. Conditional response profiles obtained from the model for different values of *τ*_1_ and *τ*_4_ (see Table 1, all other parameters as in Fig. 5) (left panel), and in the presence of mGluR_7_ receptor’s antagonist MMPIP (right panel). The latter leads to a decrease in the net amount of active mGluR_7_ and, hence, the amount of active GIRK ion channels, which corresponds to smaller values of *g_GIRK_* (see Eq. 1). Here, *τ*_1_ = 18.0s, *τ*_4_ = 120.0ms and all other parameters as in Fig. 5. Note that the normalized firing activity is calculated here by taking the inverse of the time interval between two successive spikes and dividing it by the firing frequency before the onset of the conditional response.

Changing values of both *τ*_1_ and *τ*_4_ simultaneously is one possibility to model different conditional responses within the framework of our model. We would like to point out that changing either one of the two alone does not reproduce the experimental behavior. Based on existing studies, we have no clear evidence for changes in the value of any other model parameter given our proposed mechanism. Therefore as a first approximation, we have assumed them to not change at all.

### Model predictions

Based on our proposed model, we can make two predictions that can be tested easily through experiments. 1) If PP1 is knocked out then active mGluR_7_ receptors will never deactivate after they become active from CS and hence the G-protein will remain active. This implies that the Purkinje cell will not fire again after receiving CS as shown in Fig. (8). 2) On the other hand, as PKA inhibits PP1 activity, knocking out PKA activation will activate PP1, which will dephosphorylate mGluR_7_ receptors and hence the G-protein cannot be activated. This implies that the Purkinje cell will not exhibit a conditional response as shown in Fig. (8).

**Fig 8.**
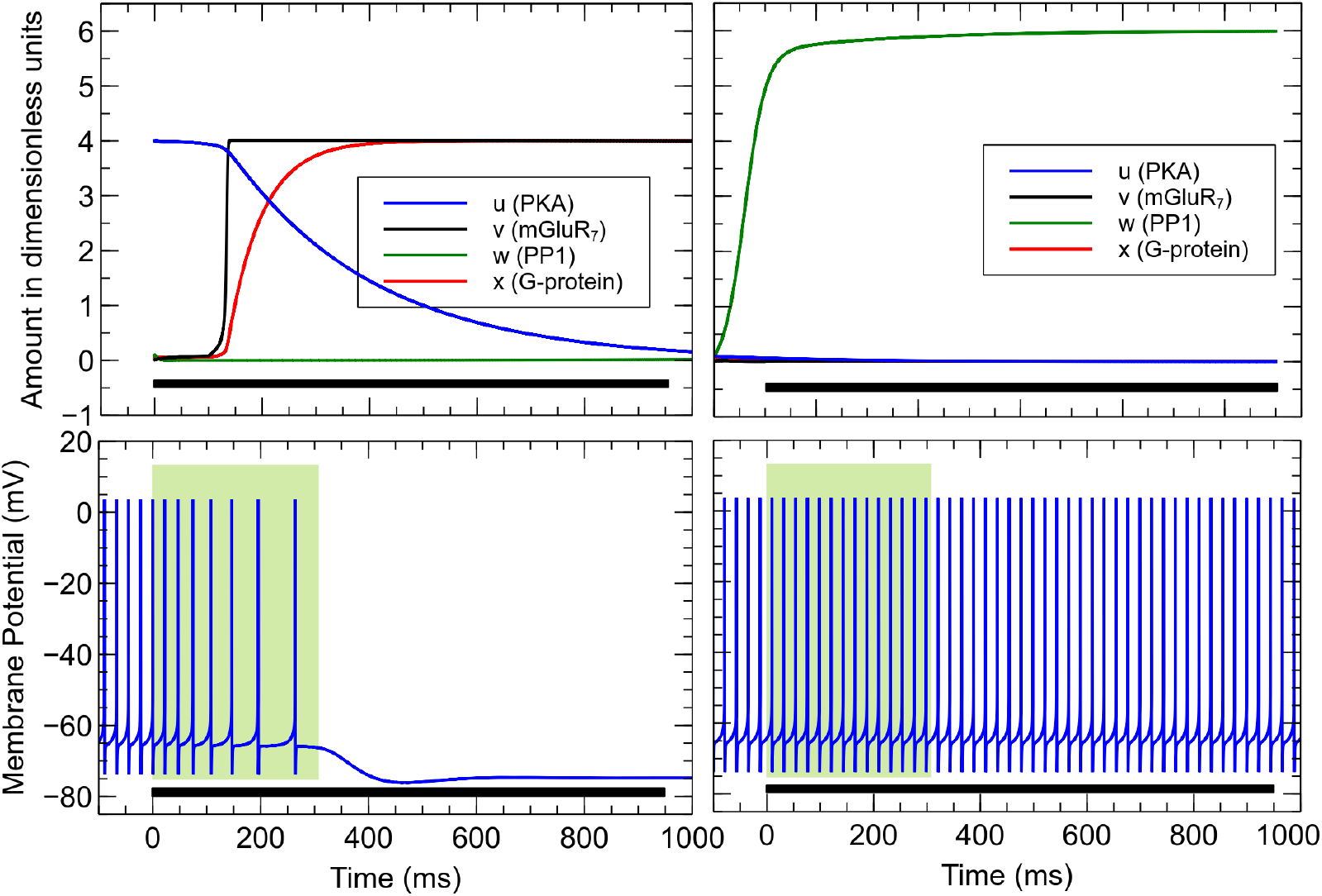
Model predictions for knockout experiments. In our mathematical model, PP1 can be knocked out by setting *w*_0_ = 0.1 (top left panel), which prevents the Purkinje cell to fire again after the initiation of the conditional response (bottom left panel). PKA can be knocked out by setting *u*_0_ = 0.1 in our model (top right panel), which prevents the Purkinje cell to initiate a conditional response (bottom right panel).

However, in reality biological cells are very robust and have redundancy mechanisms to overcome such behaviours. As a result, there might be still a weak conditional response observed after knocking out PKA or a slow deactivation of G-protein after knocking out PP1, but in both cases significant effects on the conditional response should be observed.

### Specific experimental options to test the proposed model

There are various experimental options to check whether our proposed mechanism for the conditional response is valid or not, including the two model predictions mentioned above.

1. As PP1 desensitizes the mGluR_7_ receptor during conditional response, blocking of PP1, using Okadaic acid, for example, must affect the deactivation rate of GIRK ion channels during the conditional response. This would test the first model prediction.
2. As PKA is an essential biochemical for the resensitization of the receptor and maintaining low PP1 activity, reducing PKA activity in the cell will prevent the Purkinje cell from suppressing its firing rate as PP1 will desensitize the receptor and therefore GIRK ion channels will not be activated. This can be verified by using, for example, cAMPS-Rp or triethylammonium salt, which will block the cAMP production and, hence, PKA. This would test the second model prediction.
3. If mGluR_1_ receptors are activating PKC then blocking of mGluR_1_ receptors using CPCCOEt during training will not initiate trafficking of mGluR_7_ receptors and thus no conditional response should be observed even after extensive training.
4. Use of RGS8 knockout specimen should allow only long duration conditional response: Without RGS8 protein, the activation and deactivation of G-protein will be much slower and will produce only long conditional response durations. In addition, only sufficient long CS will be able to initiate the conditional response as mGluR_7_ receptors will take longer time to activate G-protein in the absence of RGS8 protein, which accelerates G-protein activation.

## Discussion

In this article, we introduced a potential biochemical mechanism to explain time-encoding memory formation within a single synapse of a Purkinje cell. This time-encoding memory is stored in an excitatory synapse, but it is associated with an inhibitory response, i.e., the suppression of the Purkinje cell’s tonic firing rate in the presence of an excitatory stimulus, namely glutamate discharge from the parallel fiber. During conditional training, Purkinje cells imprint the time information by expressing an appropriate amount of mGluR_7_ receptors on the synapse, while encoding time information in the form of effective rate constants. The memory is stored by forming a protein complex, i.e., a Time-Encoding protein Complex (TEC). Alterations of effective rate constants within TECs will change its temporal signature, while the removal of receptors from the synapse will cause memory loss. However, during retraining, the previous memory can quickly be reacquired and it becomes accessible again. Our idea of TEC is similar to the “Timer Proteins” previously proposed by Ref. [12], but in contrast, it does not require an active selection of feedforward protein activations to produce a specific conditional response. Recently, a different biochemical mechanism was proposed for time memory learning, which uses Ca^+2^ ion dynamics for storing different time information [68]. That model does not incorporate the documented role of the GIRK ion channel and it also predicts faster learning for long duration conditional responses, which is not compatible with previous experimental findings [12].

As previously mentioned, in our model the time information of the conditional response is stored in the TECs found on individual synapses, implying that the substrate or the Engram of a time memory can reside at individual synapses, not in a cell or a cell assembly. This result is in line with the synaptogenic point of view of memory substrates [10], where single synapses play a large role in memory formation. In contrast, another point of view puts more emphasis on the intrinsic plasticity of a whole neuronal cell compared to the synaptic plasticity of individual synapses [69]. Intrinsic plasticity considers changes in the electrophysiological properties of the cell by changing the expression of Voltage-dependent Ca/K ion channels and many other kinds of ion channels, which are expressed by neurons and which decide neural firing rate as well as the sensitivity of the cell upon stimulation. However, neither points of view can fully account for the development of the conditional response in the Purkinje cell, since it neither involves the formation or elimination of pf-PC synapses [12, 13], nor LTD of pf-PC synapses [14] nor any change in the electrophysiological properties of the cell [12]. Thus, Purkinje cells show a novel form of synaptic plasticity and provide an example of monosynaptic memory encoding. In addition, considering this fact and that each Purkinje cell makes at least one synapse with up to 200,000 parallel fibers passing through the dendritic tree of the cell [70], the storage capacity of a Purkinje cell might be much higher than previously thought and the Purkinje cell might be considered as a multi-information storage device. Specifically, one might be able to encode a specific time interval by stimulating only a subset of parallel fibers and encode another time interval by stimulating a separate subset of fibers. In this case, a specific time memory out of the whole set can be selectively retrieved when the respective set of parallel fibers becomes active upon stimulation, producing the conditional response for the previously encoded time interval.

One can get a rough estimate of the total number of unique time memories that can be stored by an individual Purkinje cell by taking the ratio of the collective hyperpolarization current produced by GIRK ion channels from all synapses and the minimum required hyperpolarization current in order to noticeably suppress the tonic firing rate of the cell. To determine the minimum hyperpolarization current, one can assume that its value is approximately equal to the net resurgent Na^+^ current produced by the Purkinje cell as measured in [55]. As mentioned in the “Training with different ISI duration” subsection, the resurgent Na^+^ ion channel has the capability to spontaneously generate rapid sequences of action potentials [55, 57]. However, such resurgent currents have been observed in other neuronal classes as well, which have distinct firing activity pattern compared to the Purkinje cell [71, 72] implying that resurgent Na^+^ ion channels alone do not contribute to spontaneous high firing rates of the Purkinje cell as shown in [73]. As a result, our assumption for the minimum hyperpolarization current is just a first order approximation. In order to determine the collective hyperpolarization current, one needs to know the conductances of the GIRK ion channels and their respective densities on the synapses. Although there are experimentally measured values for single GIRK ion channel conductances [74, 75], no absolute density quantification of GIRK ion channels for Purkinje cells has been done as far as we know. Only relative abundances of GIRK ion channels at Purkinje cell’s dendritic spines are known [25]. Hence, it is currently not possible to determine the collective hyperpolarization current and, thus, the total number of unique time memories that can be stored by an individual Purkinje cell. This remains an interesting challenge for the future.

As an alternative approach, one could aim to establish experimentally that an individual Purkinje cell can indeed store at least two different time memories at separate sets of pf-PC synapses. As discussed above, stimulating separate sets of parallel fibers can in principle initiate different conditional responses. While this can be achieved by electrodes [12], it is challenging less so in terms of potential experimental protocols for conditional training [35] but rather due to difficulties in selecting specific fibers. An alternative could be to stimulate granule cells in the Granule layer of the Cerebellum [76] since parallel fibers are axonal branches of the granule cells. By stimulating a selected sub-population of granule cells and a specific branch of the climbing fiber, a subset of pf-PC synapses of an individual Purkinje cells can be trained for a specific ISI. Stimulating granule cells may appear as a drawback as they also excite other GABAergic interneurons, namely Golgi, stellate, and basket cells, which directly or indirectly can influence Purkinje cell firing activity [76]. However, their excitation did not appear to influence the conditional response profile of the Purkinje cell as shown experimentally [12]. Hence, stimulating subsets of granule cells experimentally — potentially using optogenetics [77] — might be a good way to test the capability of a Purkinje cell as a multi-information storage device in the future.

*Note added:* New support for our proposed biochemical mechanism for time-encoding memory formation comes from the observation that the mGluR_1_ receptor is necessary for the learning process, while it is not for the activation of the conditional response [78]. Despite using a different experimental approach, it basically verifies point 3 listed in the section “Specific experimental options to test the proposed model”. Specifically, the observation matches with our proposed mechanism since the latter assumes that the mGluR_1_ receptor is responsible for the learning via facilitating trafficking of the mGluR_7_ receptor to the synapses. Further experimental verification of our proposed mechanisms potentially along the lines outlined above remains an exciting challenge for the future.

## Materials and Methods

### Purkinje cell model

To model the conditional response behavior of the Purkinje cell, we start with an established dynamical model of the Purkinje cell [58] as summarized by eqs.(7) to (11). Specifically, it aims to model the dynamics of the Purkinje cell by incorporating many properties of the Purkinje cell within a realistic biophysical framework. In contrast to the original formulation [58], eqs.(7) to (11) already incorporate the features specific to our situation: In eq.(7), the input current term *I_i_*, which originally signified an external electrical stimulus, now signifies the intrinsic current causing the tonic firing of the Purkinje cell [79, 80]. Moreover, we added the influence of the GIRK ion channel in eq.(8), which only becomes relevant after training — see also eq.(1). Here, *g_GIRK_* is the net conductance of GIRK ion channels per unit area, *h_GIRK_* is the gating parameter and *V_GIRK_* is the voltage dependence of the GIRK ion channel obtained from the I-V characterstics curve of the ion channel [59].

Except for *g_GIRK_*, all values of the model are taken from [58] and listed below. As far as we know, there is no literature on the specific *g_GIRK_* values. As a result, we chose a value of *g_GIRK_* that matches the experimentally observed conditional response profiles.

Somatic voltage equation:

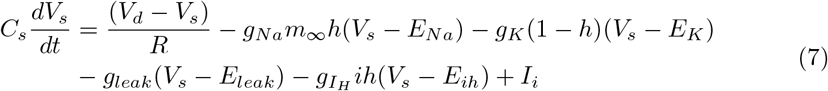

Dendritic voltage equation:

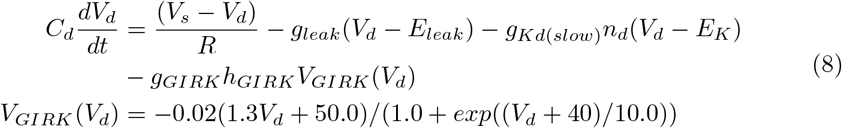

Na^+^ activation equation:

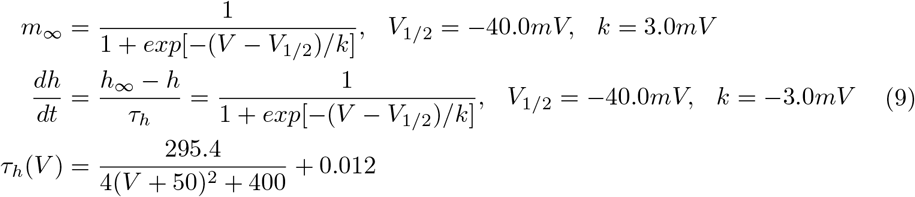

Hyperpolarizing activated cation current (I_*h*_):

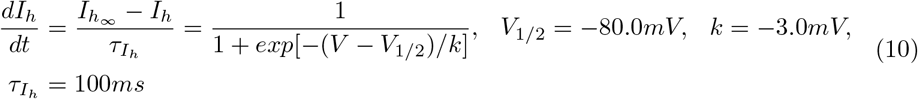

Slow K^+^ activation equation:

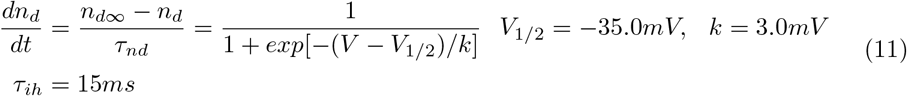

### Parameters value of the Purkinje cell model

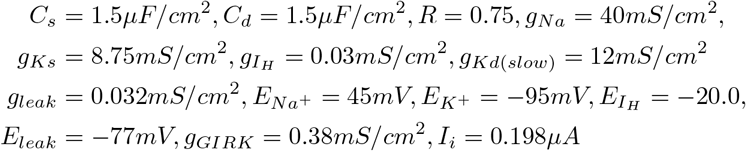

### Hill’s equation

Hill’s equation gives the fraction of protein saturated by a ligand at a given concentration of ligand in a solution [81]. Since, in the case of Acetyl cyclase (AC), only one unit of G_*α*_ subunit binds to it, Hill’s coefficient will be 1. The fraction of AC bound by a G_*α*_ subunit to the total available amount of AC at a concentration [G_*α*_] is given by

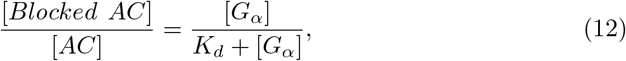

where, *K_d_* is the disassociation constant of AC and G*_α_* subunit. However, we are interested in free AC. So, the fraction of free AC will be given by

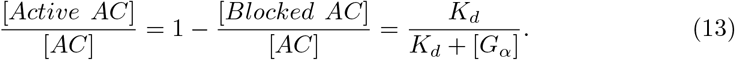

Since the concentration of [*Gα*] is proportional to its activity, eq.(13) can be written as

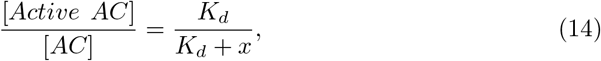

where *x* denotes the G-protein activity.

## Acknowledgments

Authors thanks Dr. Ray W. Turner at the University of Calgary for helpful discussion.

